# A haplotype-based breeding framework for the precise pyramiding of elite QTL alleles: a lettuce case study

**DOI:** 10.64898/2026.08.12.744550

**Authors:** Zhen Tu, Guangbao Luo, Li Xiao, Mingxiang Wei, Junhong Zhang, Xin Wang

## Abstract

The efficient pyramiding of favorable alleles underlying complex traits remains a major challenge in crop breeding as most quantitative trait loci (QTLs) have not been resolved to causal genes, limiting their direct application in marker-assisted breeding. Although haplotypes provide more informative genetic units than individual markers, existing haplotype-based studies have largely focused on genetic interpretation and elite haplotype discovery, whereas computational frameworks for translating haplotypes into breeding decisions remain limited. Here, we developed HAPBDB, a haplotype-guided breeding framework that directly translates regional haplotypes into parental selection, cross design, and elite QTL pyramiding, and applied it to a lettuce genomic breeding panel. HAPBDB accurately reconstructed functional haplotypes at known loci and resolved elite haplotypes for five major QTLs controlling flowering time and yield. Integrating haplotype information across loci enabled systematic identification of accessions carrying complementary elite haplotypes and rational design of crosses that maximized favorable haplotype accumulation while minimizing segregating loci. Experimental validation using QTL-specific molecular markers demonstrated concordance between predicted and observed multi-locus genotypes across all designed F hybrids. Our results demonstrated that regional haplotypes can serve as practical breeding units even when the underlying causal genes remain unknown, thereby enabling the direct utilization of genetically mapped QTLs for precision breeding. By bridging the gap between genomic discovery and practical breeding, HAPBDB provides a practical framework for converting genomic information into breeding decisions and accelerating precision improvement of complex traits.

## Introduction

Crop breeding aims to develop improved varieties with higher yield, enhanced stress tolerance, and better nutritional quality to meet the growing demands of global agriculture ^1,2^. With the advent of Mendelian genetics and modern molecular biology, breeding practices have evolved toward more targeted and predictive approaches. Marker-assisted selection (MAS), which uses molecular markers linked to favorable alleles, has greatly accelerated crop improvement ^3,4^. However, most agronomically important traits, including yield, flowering time, grain quality, and stress tolerance, are quantitatively inherited and controlled by numerous loci with small individual effects ^5,6^. Consequently, a single marker or gene cannot fully explain the variation of these traits, and selection based on individual loci only captures part of the genetic variance. Accordingly, the integration of elite alleles from multiple loci into a single genetic background has become a central objective of modern breeding, a process commonly referred to as allele pyramiding. However, pyramiding multiple favorable loci into a single cultivar often requires many years of repeated crossing, selection, and phenotypic evaluation, making the process labor-intensive, time-consuming, and inefficient ^7,8^. Rational breeding strategies that enable the simultaneous aggregation of multiple elite alleles through genomics-guided parental selection could rapidly generate individual plants carrying the desired allele combinations in one or two crosses, escaping multiple rounds of phenotypic evaluation and selection ^9,10^. Such strategies can substantially shorten breeding cycles while reducing labor and resource requirements by enabling early-generation parental selection and marker-assisted breeding.

Recent advances in high-throughput genotyping and phenotyping technologies have greatly improved our ability to dissect the genetic architecture of complex traits at increasingly higher resolution ^11^. Genome-wide association studies (GWAS) and bulked segregant analysis (BSA) have become powerful approaches for identifying quantitative trait loci (QTLs) associated with agronomic traits ^4,12,13^. These approaches enable the detection of both major- and minor-effect loci across the genome, providing a comprehensive view of the genetic basis of complex traits. However, identifying the causal genes underlying these QTLs remains challenging because the extent of linkage disequilibrium (LD) often limits mapping resolution ^14^ In many crop species, particularly those that have undergone recent strong domestication bottlenecks, intense artificial selection, or clonal propagation, LD could extend over megabase scale genomic regions, resulting in even hundreds of genes are inherited as a single haplotype block ^6,15^. Consequently, although GWAS and BSA are powerful approaches for detecting trait-associated regions, fine mapping of causal genes within these intervals is often prohibitively time-consuming and may be impractical in genomic regions with severely suppressed recombination, such as chromosomal inversions ^2^, pericentromeric regions ^16^ and telomeric regions 17. Therefore, for many breeding applications, the trait-associated haplotype itself rather than the causal gene may represent the most practical unit for marker-assisted selection and molecular breeding.

Compared with individual SNP markers, haplotypes represent combinations of linked variants that are inherited together and therefore capture a broader spectrum of genetic variation. Because they integrate the effects of multiple co-inherited variants, haplotypes provide a more robust representation of functional genetic diversity, particularly in genomic regions with strong LD or limited recombination ^18^. In addition, the multi-allelic nature of haplotypes makes them inherently more informative than biallelic SNP markers for both genetic analysis and breeding ^19^. Accordingly, haplotype-based analyses have become increasingly important for dissecting the genetic basis of complex traits and numerous studies have shown that haplotype-based GWAS could achieve higher mapping power and accuracy than conventional SNP-based analyses ^20,21^. More importantly, haplotypes provide practical breeding units because they capture the combined effects of linked functional variants even when the underlying causal mutations remain unknown. In recent years, haplotype-based breeding has emerged as a powerful strategy in crop breeding by analyzing haplotype blocks instead of individual SNPs to improve breeding efficiency and genetic gain ^22^. In rice, elite haplotypes identified from the 3K Rice Genomes Project have been successfully exploited to improve grain yield and quality without prior identification of the causal variants ^18^. Similar approaches have also been applied in pigeonpea to identify elite haplotypes associated with drought tolerance ^23^, demonstrating the broad applicability of haplotype-assisted breeding across crop species. These studies highlight that haplotypes constitute a more effective selection unit than individual markers for improving complex traits. Despite these advances, current studies have largely focused on identifying superior haplotypes associated with target traits ^9,22^. Computational frameworks that systematically integrate multiple elite haplotypes, optimize parental selection, and rationally design breeding combinations remain largely unavailable.

To address this gap, we developed HAPBDB, a computational toolkit for regional haplotype reconstruction, clustering, visualization, and multi-locus genotype analysis using large-scale genomic variation datasets. Using a lettuce genomic breeding panel, we reconstructed regional haplotypes for five major QTLs controlling flowering time and yield, identified elite haplotypes, and integrated them across loci to design optimal breeding combinations. Molecular validation using QTL-specific InDel markers confirmed the accuracy of the predicted multi-locus genotypes in designed hybrids. Collectively, our results demonstrate that regional haplotypes can serve as actionable breeding units for parental selection, cross design, and elite QTL pyramiding, providing a practical framework for accelerating precision breeding in crop species.

## Results

### Overview of the HAPBDB (Haplotype-Based Design Breeding) workflow

To facilitate the efficient utilization of elite QTLs in molecular breeding, we developed HAPBDB that integrates haplotype identification, multi-locus genotype analysis, and breeding design into a unified analytical framework (**Fig. 1**). The workflow consists of four sequential modules: (i) construction of a high-confidence variant matrix, (ii) haplotype identification and clustering, (iii) multi-QTL haplotype integration, and (iv) breeding design for elite haplotype pyramiding.

**Figure 1.**
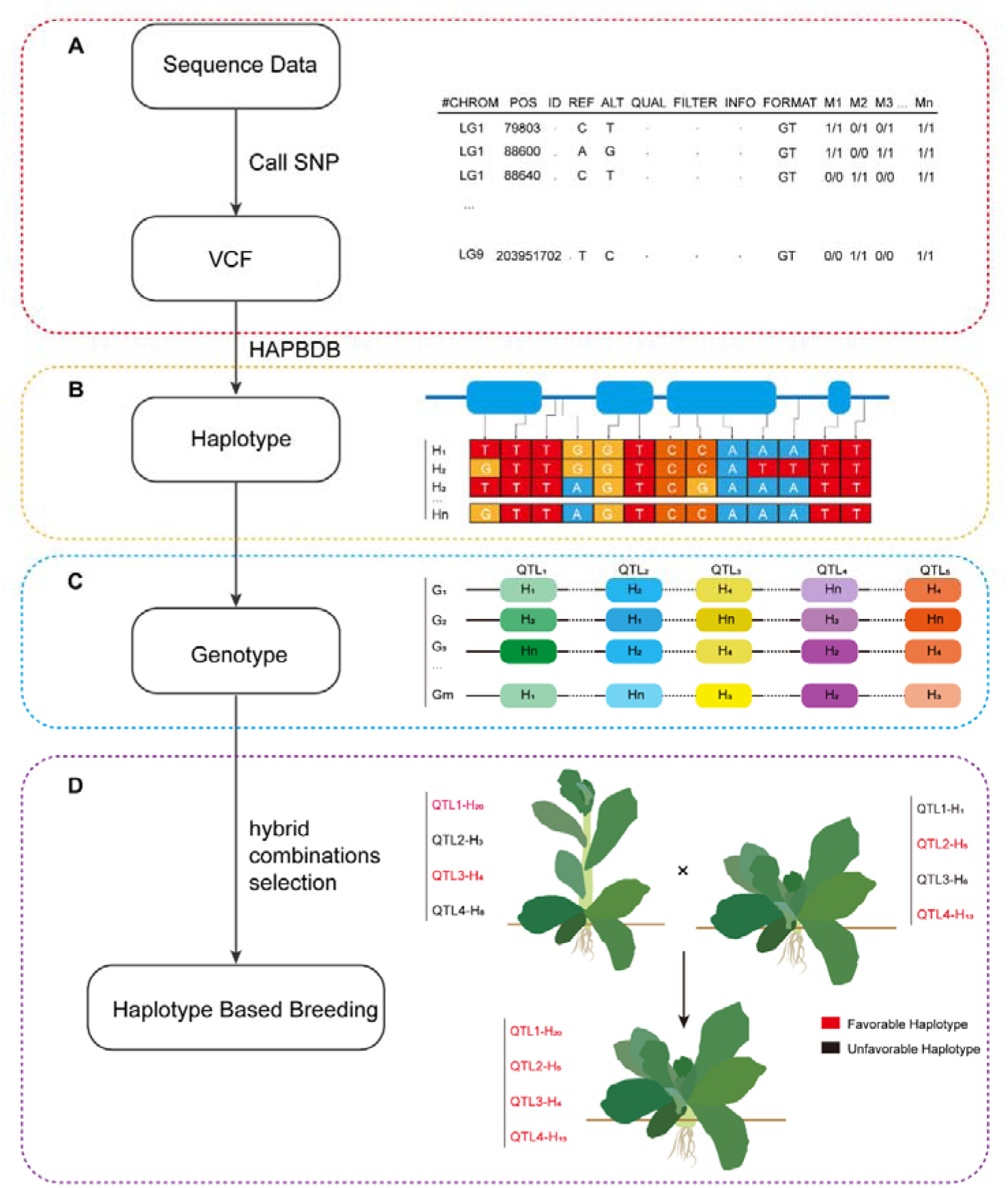
Overview of HAPBDB workflow. The workflow consists of four sequential modules (left), with representative outputs illustrated on the right. (**A**) SNP calling and construction of a high-confidence VCF file using GATK. The red panel shows an example of the resulting VCF format. (**B**) Regional haplotype reconstruction and clustering using HAPBDB. The yellow panel illustrates representative haplotypes (H1–Hn) identified within a target genomic region. (**C**) Multi-locus haplotype integration across target QTLs. The blue panel shows the combinations of haplotypes across five QTL regions, generating distinct multi-locus genotypes (G1–Gm). (**D**) Haplotype-guided breeding design. The purple panel illustrates the selection of complementary parental combinations to pyramid elite haplotypes from multiple QTLs into a single breeding line.

The workflow begins with a high-quality genomic variant matrix generated from whole-genome resequencing data. Following standard variant calling and quality filtering, reliable SNPs and InDels within target QTL intervals are extracted to construct the genotype matrix used for downstream haplotype analysis (**Fig. 1A**).

The filtered variant matrix is subsequently analyzed using HAPBDB, a Python-based toolkit specifically developed for haplotype analysis within trait-associated genomic regions. HAPBDB employs a tree-guided hierarchical clustering strategy that groups genetically similar accessions according to sequence similarity while separating divergent haplotypes based on pairwise genetic distances. In addition to haplotype classification, HAPBDB generates intuitive visualizations, including haplotype dendrograms, Hamming distance heatmaps, and base-level haplotype matrices, allowing users to examine sequence variation and genetic relationships among haplotypes within individual QTL regions (**Fig. 1B**).

Following haplotype reconstruction, each accession is assigned to a specific haplotype at every target QTL. Using user-defined elite and non-elite reference haplotypes, HAPBDB automatically classifies each accession and integrates haplotype information across all selected QTLs to generate a multi-locus haplotype matrix. This matrix summarizes the distribution of elite haplotypes within the breeding population and enables rapid identification of accessions carrying complementary favorable haplotypes across multiple loci (**Fig. 1C**).

The integrated haplotype matrix subsequently serves as the basis for breeding design. Candidate parental lines are prioritized according to the complementarity of elite haplotypes, allowing breeders to select crosses that maximize the accumulation of favorable alleles in the progeny. Rather than evaluating individual QTLs independently, the workflow simultaneously considers all target loci and predicts the haplotype composition expected in each cross, thereby facilitating the rational design of parental combinations for multi-QTL pyramiding (**Fig. 1D**).

Collectively, HAPBDB establishes an end-to-end framework that links genomic variation, regional haplotype analysis, and breeding design within a single pipeline, transforming them into actionable units for parental selection, cross design, and elite haplotype pyramiding. By integrating haplotype reconstruction, multi-locus genotype analysis, and breeding optimization, HAPBDB provides a practical strategy for accelerating precision breeding of complex traits.

### Development of a representative lettuce genomic breeding panel for haplotype-guided analysis

To establish a comprehensive genomic resource for genetic diversity analysis and precision molecular breeding in lettuce, we constructed a high-resolution lettuce genomic breeding (LG) panel by integrating large-scale variation datasets derived from both whole-genome resequencing (WGS) and transcriptome sequencing. Specifically, the panel combines genomic variation data from a MAGIC (multi-parent advanced generation intercross) population and a natural population, thereby capturing complementary sources of allelic diversity. A total of 46,217,729 SNPs were initially identified from the MAGIC population WGS dataset, while 401,272 SNPs were obtained from transcriptome sequencing of the natural population. After intersecting and harmonizing the datasets, 203,290 high-confidence SNP overlapped loci were retained for panel construction. These shared SNPs provided a unified variant backbone for downstream population characterization and haplotype analysis. The resulting LG panel comprised 631 accessions, representing the major cultivated types of *Lactuca sativa* as well as its closely related wild species (**Table S1**). The cultivated germplasm included 28 Butterhead, 21 crisphead (head), 17 looseleaf, 31 romaine (cos type), 33 stem lettuce, and 379 MAGIC. In addition, wild relatives were incorporated to broaden the genetic base, including *L. serriola* (n = 27), *L. saligna* (n = 2), and *L. virosa* (n = 2). These accessions were collected from major lettuce production regions across Asia, Europe, and North America, as well as from international germplasm repositories. This broad geographic and taxonomic sampling enabled the LG panel to capture both domestication related diversity and wild allelic variation.

To evaluate whether the LG panel represented the genetic diversity of previously characterized lettuce germplasm, we performed comprehensive population genetic analyses of LG panel together with previously published lettuce variation map comprising 445 accessions. Phylogenetic analysis of 1,076 accessions revealed that the LG panel accessions were distributed across all major clades previously described in cultivated and wild lettuce (**Fig. 2A**). Principal component analysis (PCA) produced a population structure consistent with the phylogenetic relationships (**Fig. 2B**), indicating that the LG panel retained the major genetic lineages present in global lettuce diversity. Population structure analysis at K = 5 revealed clear differentiation among major horticultural groups, almost consistent with the phylogenetic tree of LG panel (**Fig. 2C**). Notably, accessions derived from the MAGIC population were positioned between wild relatives and cultivated lettuce, suggesting that these lines harbor recombined haplotypes originating from diverse parental backgrounds. This intermediate genetic position indicated that MAGIC-derived lines may contain multiple favorable alleles within a single accession, making them particularly valuable as bridging or donor parents in breeding programs. Collectively, the LG panel represents a genetically diverse and well-characterized genomic resource that integrates cultivated germplasm, wild relatives, and recombined breeding materials, providing a robust foundation for regional haplotype reconstruction, elite haplotype identification, and haplotype-guided breeding design in lettuce.

**Figure 2.**
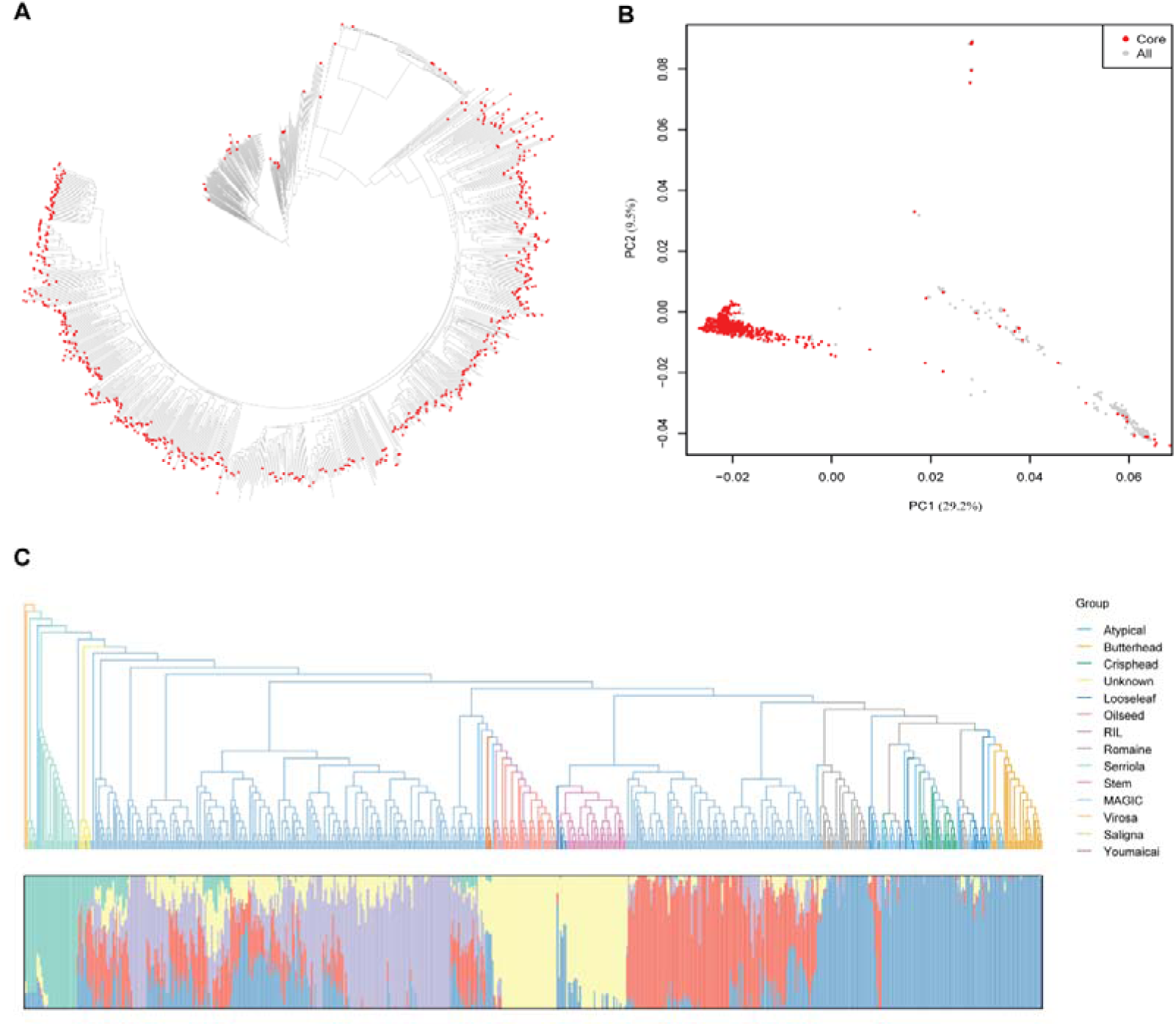
Genetic characterization of the lettuce genomic breeding (LG) panel. (**A**) Phylogenetic analysis of 1,076 lettuce accessions. Accessions included in the LG panel are highlighted in red. (**B**) Principal component analysis (PCA) of the same accessions, with LG panel accessions shown in red. (**C**) Phylogenetic tree and model-based population structure (K = 5) of the LG panel.

### HAPBDB accurately reconstructs functional regional haplotypes at known loci

To evaluate the accuracy of HAPBDB in reconstructing regional functional haplotype, we first examined several previously characterized functional loci within the LG panel. As a representative example, we analyzed the *LsphyB* locus, in which a nonsense mutation has previously been reported in the late-flowering cultivar Wo111 ^12^. This causal mutation results corresponds to a G-to-A substitution at position 47,654,275 on chromosome 1, causing premature termination of the LsphyB protein. Variants within the surrounding genomic interval (Chr1: 47,337,800–47,930,000) that represented the putative QTL of *LsphyB* were extracted from the LG panel and subjected to haplotype reconstruction using HAPBDB (**Fig. 3A**).

**Figure 3.**
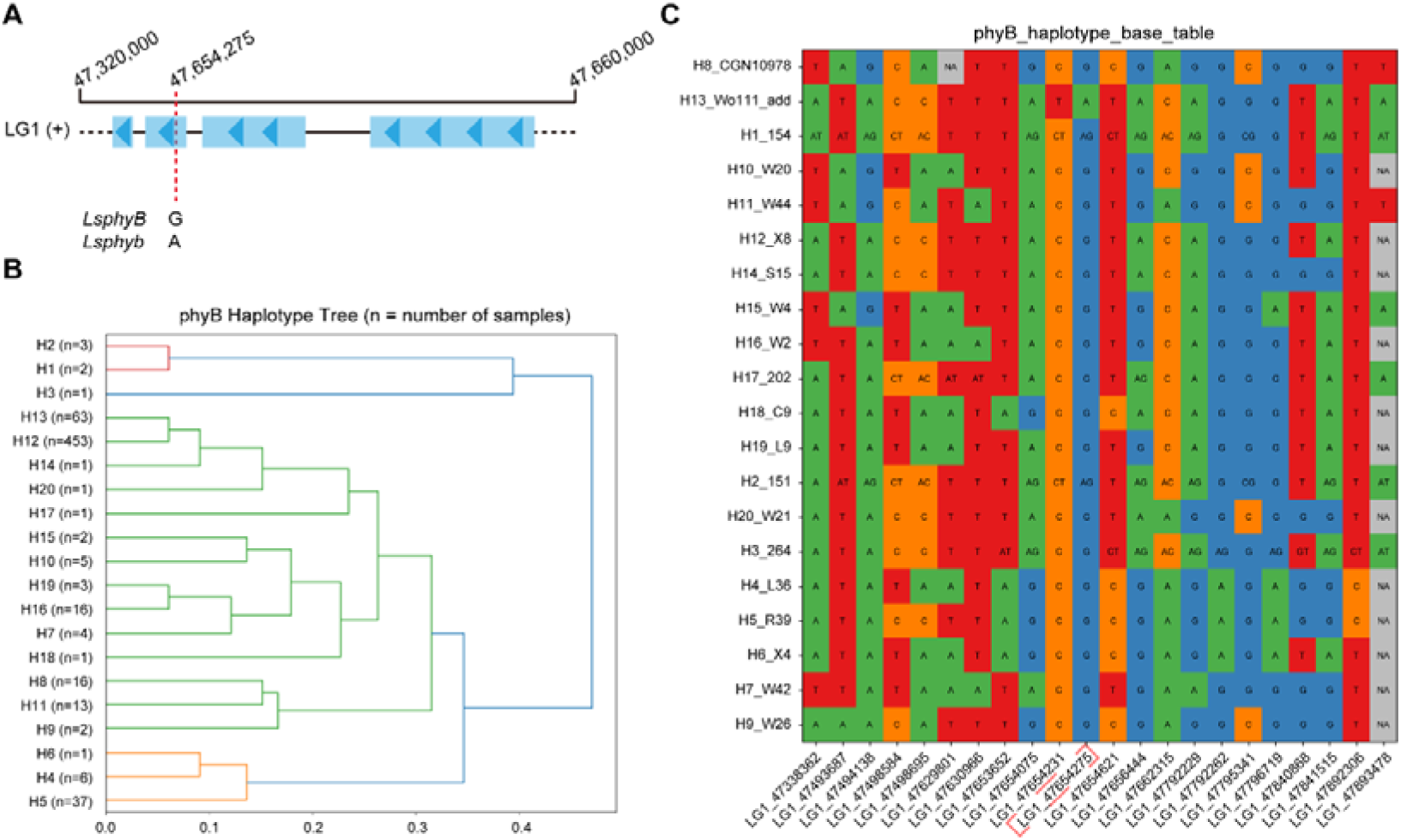
Regional haplotype dissection of the *LsphyB* locus using HAPBDB. (**A**) Genomic organization of the *LsphyB* locus (LG1: 47,320,000–47,660,000), with the reported causal SNP highlighted in red box. (**B**) Dendrogram showing the relationships among regional haplotypes reconstructed by HAPBDB. Branch lengths are proportional to pairwise Hamming distances, with closely clustered haplotypes exhibiting higher sequence similarity. (**C**) Haplotype matrix showing nucleotide variation across representative regional haplotypes. Haplotypes are grouped according to their phenotypic effects (unfavorable, favorable, and others), facilitating the identification of sequence patterns associated with the favorable allele.

HAPBDB classified this genomic region into 21 distinct haplotypes, with H12 representing the predominant haplotype carried by 453 accessions (**Fig. 3B**). The late-flowering accession Wo111 was assigned to haplotype H13, which comprised 63 accessions. In contrast, all remaining haplotypes contained either the reference G allele or the heterozygous A/G genotype at this position (**Fig. 3C**). Notably, the dominant H12 haplotype was present in predominantly cultivated accessions, suggesting that this genomic region has been shaped during lettuce domestication and breeding.

To further assess the general applicability of HAPBDB, two additional loci controlling leaf angle, *LsKIPK* and *LsATPase*, were analyzed using the same workflow (Xie et al., 2024). At the *LsKIPK* locus, the previously reported causal mutation was assigned to haplotype H1, with 249 of 252 accessions (98.8%) showing concordant classification with the reported genotypes (**Fig. S1**). Similarly, the haplotype harboring the causal mutation of *LsATPase* was assigned in H9, correctly classifying 75 of 78 accessions (96.2%) (**Fig. S2**). These results demonstrated that HAPBDB accurately reconstructs regional haplotypes across diverse genomic loci and effectively groups accessions carrying identical functional alleles. More importantly, this feature makes HAPBDB particularly valuable for molecular breeding, where many target QTLs have been genetically mapped but their underlying causal genes remain unresolved.

### Identification of elite regional haplotypes across breeding-relevant QTLs

To demonstrate the application of HAPBDB in genomic regions harboring the breeding allele, we performed haplotype analyses for five previously identified QTLs controlling flowering time and yield in lettuce. These included two flowering-time QTLs (FT_Wo46 and FT_Y37) and three yield-related QTLs (YD_Wo328, YD_Y37 ^24^, and YD_Wo47), each originating from elite donor accessions identified through genetic mapping. For each QTL interval, regional sequence variation across the LG panel was extracted and analyzed using HAPBDB to reconstruct haplotype structures and identify donor-derived elite haplotypes (Table S2). Following our nomenclature of their donor origins, elite flowering-time and yield haplotypes were designated as FT_x_Hy and YD_x_Hy, respectively, where x denotes the donor accession from which the elite haplotype was originally identified and y represents the elite haplotype group. Distinct haplotype structures were successfully resolved for all five QTLs, allowing the elite donor haplotypes to be unambiguously distinguished from alternative haplotypes (**Fig. 4A, B**). Specifically, HAPBDB identified extensive haplotype diversity across these QTL regions, resolving 50 haplotypes at FT_Wo46, 15 at FT_Y37, 21 at YD_Wo328, 136 at YD_Y37, and 358 at YD_Wo47. The elite haplotypes carried by the donor parents corresponded to FT_Wo46_H4, FT_Y37_H4, YD_Wo328_H3, YD_Y37_H21, and YD_Wo47_H5, respectively, demonstrating that regional haplotype reconstruction enables the identification of favorable genetic configurations within highly diverse QTL intervals.

**Figure 4.**
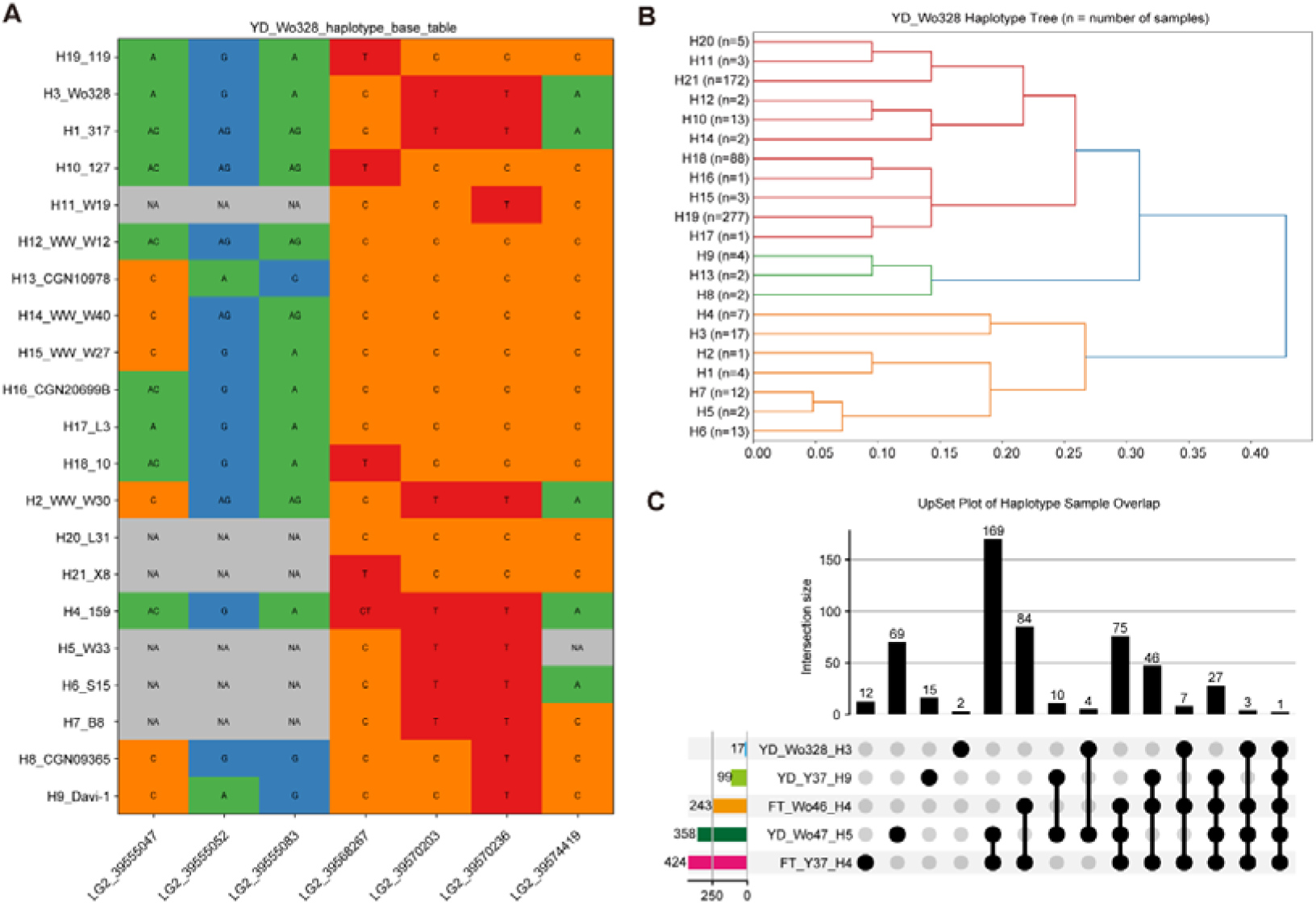
Identification and integration of elite regional haplotypes across five target QTLs. (**A**) Genomic organization and regional haplotype reconstruction of the representative yield QTL YD_Wo328. (**B**) Dendrogram illustrating the relationships among regional haplotypes reconstructed by HAPBDB, with branch lengths proportional to pairwise Hamming distances. (**C**) UpSet plot summarizing the combinations of elite haplotypes across the five target QTLs. Bars represent the number of accessions carrying each haplotype combination, and connected dots indicate the corresponding QTLs.

We next investigated the distribution and accumulation of these elite haplotypes across the LG panel. The analysis revealed substantial variation in the accumulation of favorable haplotypes among accessions (**Table S3**). A total of 169, 84, 10, and 4 accessions carried elite haplotypes at two target QTLs. Notably, 169 accessions simultaneously harbored the elite haplotypes of YD_Wo47 and FT_Y37, suggesting that these elite haplotypes have been widely incorporated into lettuce breeding germplasm. In addition, 74, 46, and 7 accessions carried elite haplotypes at three QTLs, whereas 27, 3, and 1 accessions possessed favorable haplotypes at four QTLs. Most notably, one MAGIC-derived accession (260) contained elite haplotypes at all five target loci, representing an ideal donor for multiple traits improvement through either direct selection or strategic hybridization (**Fig. 4C**). Together, these analyses demonstrated that HAPBDB not only accurately reconstructs regional haplotypes but also enables systematic evaluation of elite haplotype combinations across breeding populations. By transforming QTL related variation into interpretable haplotype units, HAPBDB provides a practical basis for identifying superior parental materials and designing crosses for multi-QTL pyramiding.

### Haplotype-guided breeding design identifies optimal parental combinations for elite haplotype pyramiding

To demonstrate the practical application of HAPBDB in breeding design, we next evaluated multilocus haplotype configurations across the LG panel to identify parental combinations capable of maximizing elite haplotype accumulation of elite haplotypes across the five target QTLs. Based on the haplotype profiles generated by HAPBDB, six accessions carrying complementary elite haplotypes were selected as candidate parents for breeding (**Fig. 5A**). Among the five designed crosses, accessions 265 and 324 each possessed four elite haplotypes, whereas accessions 159 and Wo328 carried three, and accessions 119 and 375 contained two elite haplotypes. Together, these accessions represented an optimal parental pool for designing multi-trait breeding combinations. Using these six accessions, five hybrid combinations were designed to maximize the number of elite haplotypes transmitted to the progeny (**Fig. 5B; Table S4**). Three combinations (1_F1, 2_F1, and 3_F1) were predicted to integrate all five elite haplotypes into the F generation, whereas two additional combinations (4_F1 and 5_F1) were expected to combine four elite haplotypes. Depending on the parental haplotype composition, elite haplotypes were predicted to be either homozygous or heterozygous in the resulting hybrids, thereby determining the probability of obtaining completely fixed elite genotypes in subsequent generations.

**Figure 5.**
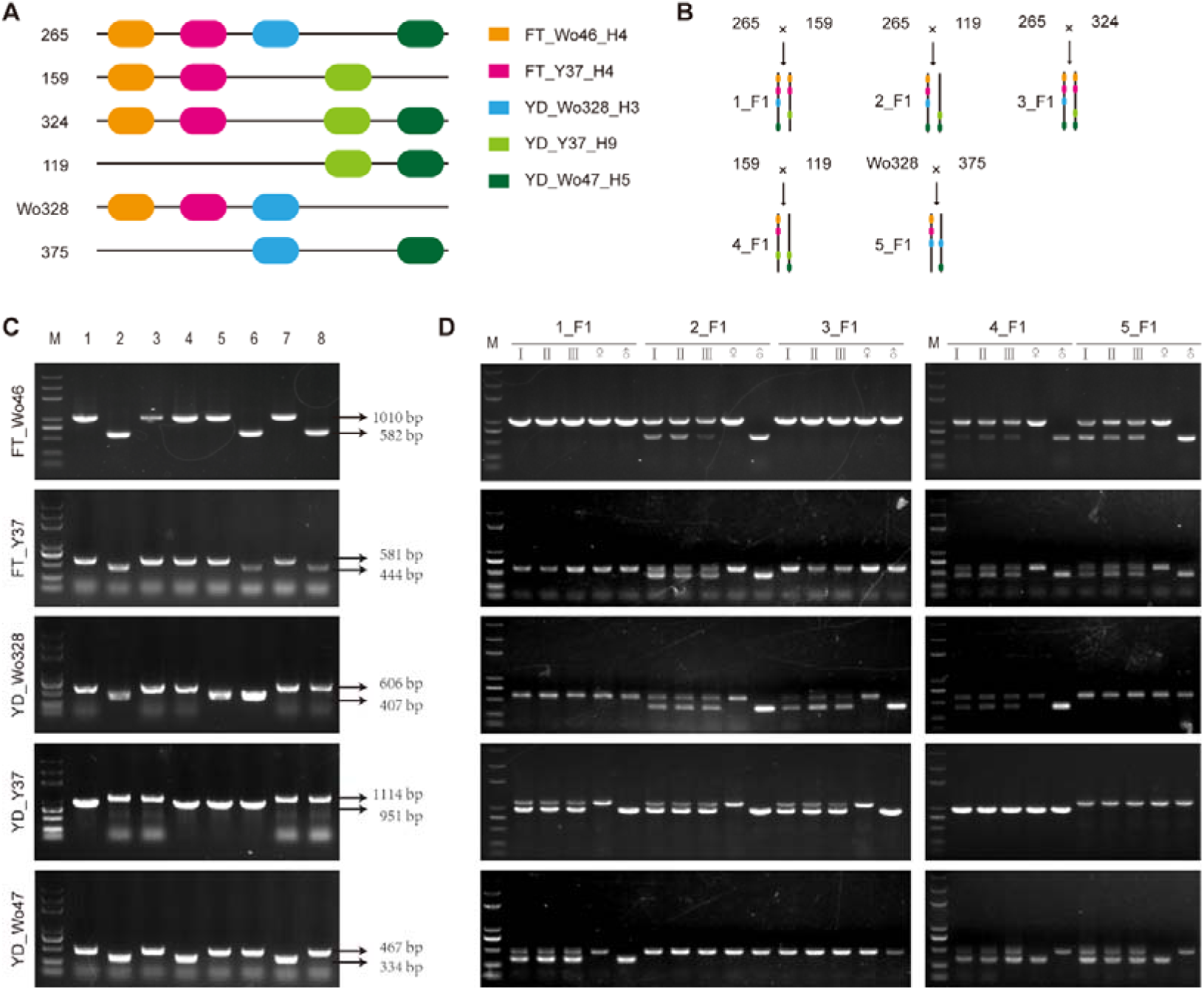
Haplotype-guided breeding design and validation of elite haplotype pyramiding. **(A)** Distribution of elite regional haplotypes across the five target QTLs in six selected parental accessions. **(B)** Rational design of five hybrid combinations to maximize the pyramiding of elite regional haplotypes. Colored connections indicate the parental origin of each elite haplotype, and the predicted number of elite haplotypes in each F hybrid is shown. **(C)** Representative validation of parental haplotypes using QTL-specific InDel markers. Lanes 1 and 2 correspond to the original QTL mapping parents carrying the elite and non-elite haplotypes, respectively, whereas lanes 3–8 represent the six selected parental accessions. **(D)** Experimental validation of the predicted haplotype composition in the five designed F hybrid combinations using the corresponding InDel markers. Symbols ♀ and ♂ indicate the female and male parents, respectively, and I–III represent three independent F plants from each cross.

Among the five designs, 3_F1 (265 × 324) represented the most favorable breeding combination. Because the two parents shared three elite haplotypes while contributing complementary elite haplotypes at the remaining two loci, the resulting F was predicted to contain three homozygous and two heterozygous elite loci. Under Mendelian segregation, only the two heterozygous loci require fixation in the F generation, resulting in a theoretical recovery frequency of 1/16 (6.25%) for individuals homozygous for all five elite haplotypes, indicating only one generation of selfing would be required to recover completely homozygous elite genotypes at all target QTLs. In contrast, breeding designs containing three or four heterozygous loci would produce substantially lower recovery frequencies (1/64 and 1/256, respectively), illustrating the importance of rational parental selection for accelerating elite haplotype pyramiding. These results demonstrate that HAPBDB not only identifies elite haplotypes but also provides a quantitative framework for optimizing parental selection and predicting the haplotype composition of progeny. Rather than relying on empirical crossing strategies, the workflow enables breeders to rationally design crosses that maximize elite haplotype accumulation while minimizing the number of segregating loci, thereby substantially improving breeding efficiency.

### Experimental validation confirms the accuracy of haplotype-guided breeding predictions

To experimentally validate the haplotype-assisted breeding designs generated by HAPBDB, we developed InDel markers for each of the five target QTLs and genotyped the selected parental accessions and their corresponding F hybrids. Marker-assisted genotyping was used to evaluate whether the predicted haplotype composition generated by HAPBDB accurately reflected the genetic constitution of the breeding materials. We first examined polymorphisms among the six selected parental accessions. PCR amplification showed that the marker genotypes were highly consistent with the haplotype classifications generated by HAPBDB across all five target QTLs (**Fig. 5C**; **Table 1**). A single exception was observed at YD_Wo328 in accession 159, where the InDel marker detected the elite genotype, whereas HAPBDB assigned the accession to haplotype YD_Wo328_H4 rather than the elite haplotype YD_Wo328_H3. Inspection of the regional haplotype structure revealed that H3 and H4 formed adjacent branches in the phylogenetic tree and differed only at two heterozygous SNPs while sharing otherwise identical sequence composition (**Fig. 4A, B**). This result indicated that HAPBDB distinguishes closely related regional haplotypes based on complete sequence variation rather than a single marker, highlighting its higher resolution for regional haplotype characterization. Importantly, this minor discrepancy did not affect the identification of the elite haplotype or subsequent breeding design.

**Table 1.** Correspondence between parental QTL haplotypes and InDel markers.

| Parent | 265 |  | 159 |  | 324 |  | 119 |  | Wo328 |  | 375 |  |
| --- | --- | --- | --- | --- | --- | --- | --- | --- | --- | --- | --- | --- |
|  | Hap | Ind | Hap | Ind | Hap | Ind | Hap | Ind | Hap | Ind | Hap | Ind |
| FT_Wo46 | + | + | + | + | + | + | - | - | + | + | - | - |
| FT_Y37 | + | + | + | + | + | + | - | - | + | + | - | - |
| YD_Wo328 | + | + | - | + | - | - | - | - | + | + | + | + |
| YD_Y37 | - | - | + | + | + | + | + | + | - | - | - | - |
| YD_Wo47 | + | + | - | - | + | + | + | + | - | - | + | + |

+: elite haplotype; -: non-elite haplotype; Hap: haplotype; Ind: indel marker.

The same set of InDel markers was subsequently applied to genotype all five designed F hybrids. In every hybrid combination, the observed marker genotypes matched the haplotype compositions predicted by HAPBDB (**Fig. 5D; Table S5**). Homozygous and heterozygous elite haplotypes at each QTL were detected exactly as predicted from the parental haplotype compositions, demonstrating that HAPBDB accurately predicts multi-locus genotype composition in designed crosses. Collectively, these experimental results provided molecular validation of the HAPBDB breeding framework. The close agreement between predicted and observed multiple locus genotypes demonstrated that regional haplotypes can be reliably translated into practical breeding decisions, thereby establishing HAPBDB as an effective framework linking genomic variation with precision breeding.

## Discussion

### Advances of Haplotype-Based Breeding

Improving complex agronomic traits remains a major challenge in crop breeding because these traits are typically controlled by numerous loci with small individual effects ^13,25^. Although marker-assisted selection (MAS) and genomic selection (GS) have substantially accelerated crop improvement, both approaches have inherent limitations ^3,10^. MAS generally relies on individual markers within QTL intervals, whose predictive power may be compromised by recombination or incomplete linkage with causal variants. In contrast, GS captures genome-wide marker effects to achieve high predictive accuracy but provides limited biological interpretation and little guidance for targeted allele pyramiding. Haplotypes, which integrate multiple linked variants inherited together, provide a more informative representation of functional genetic variation by capturing additive, dominant, and epistatic effects that are often overlooked by single-marker analyses ^26^. Consequently, haplotype-based breeding has emerged as a promising strategy for improving complex traits. However, most existing studies have focused on haplotype discovery, population genetics, or association analyses, with relatively few approaches translating haplotype information into practical breeding decisions ^20–22^.

In this study, we established a haplotype-assisted breeding framework that directly links QTL information to breeding design. Instead of relying on individual diagnostic markers, our approach reconstructs regional haplotypes within target QTL intervals and identifies elite haplotypes associated with desirable agronomic traits. These haplotypes are subsequently integrated across multiple QTLs to evaluate breeding materials, identify complementary parental lines, and design crosses that maximize elite haplotype pyramiding. Importantly, this strategy does not require prior identification of the causal gene, making it particularly valuable for many breeding-relevant QTLs that have been genetically mapped but remain molecularly unresolved. Rather than using haplotypes solely to interpret genetic variation, our framework establishes haplotypes as actionable units for parental selection, cross design, and elite haplotype pyramiding, thereby bridging the gap between genomic discovery and practical molecular breeding.

### Application of HAPBDB in polygenic trait improvement

The successful pyramiding of elite haplotypes in lettuce demonstrates the practical utility of haplotype-assisted breeding for improving complex traits. More broadly, the proposed framework is readily applicable to other crop species in which large-scale resequencing datasets and genetically mapped QTLs are available, even when the underlying causal genes remain unknown. Rather than treating individual QTLs independently, the framework evaluates complementary haplotype combinations across multiple loci simultaneously, enabling rational selection of parental lines and optimization of breeding strategies.

This approach is particularly well suited for polygenic traits such as yield, flowering time, quality, and abiotic or biotic stress tolerance, which are typically controlled by multiple loci with moderate or small effects. In crops such as rice and maize, where extensive haplotype resources have already been established ^18,23^, the framework could complement genomic selection by providing biologically interpretable selection units and facilitating targeted elite haplotype pyramiding. Consequently, HAPBDB extends the application of haplotype analysis from genetic discovery to practical molecular breeding and provides a general strategy for design-based crop improvement.

### Limitations and future perspectives

Despite its advantages, several challenges remain for haplotype-assisted breeding. First, the effectiveness of regional haplotypes is strongly influenced by local linkage disequilibrium (LD) patterns, which differ among species and populations ^10^. In genomic regions with severely suppressed recombination, such as chromosomal inversions and pericentromeric regions, large haplotype blocks may contain both favorable and unfavorable alleles, increasing the risk of linkage drag and reducing breeding efficiency. Emerging genome engineering technologies, including CRISPR-based chromosome engineering, may provide opportunities to overcome these constraints by breaking unfavorable linkage relationships ^27,28^. Second, although the current study developed and validated HAPBDB using diploid lettuce populations, the underlying analytical framework has subsequently been extended to support polyploid genomes through allele-dosage-aware genotype encoding and phased haplotype representation, allowing both diploid and polyploid datasets to be processed within a unified analytical framework. This extension enables the incorporation of allele dosage and haplotype information into a unified analytical workflow, broadening the potential applicability of HAPBDB to polyploid crops. However, the comprehensive evaluation of the extended framework in polyploid species remains necessary, particularly for accurate haplotype inference, homologous chromosome phasing, and the assessment of breeding performance across different ploidy levels. Finally, breeding decisions based solely on haplotype composition may not fully capture the effects of genotype-by-environment interactions or epistatic interactions among loci ^29,30^. Integrating haplotype-based breeding design with genomic prediction, multi-environment phenotyping, and functional genomic information will further improve the robustness and predictive power of the framework. Such integration will facilitate the development of superior cultivars with stable performance across diverse genetic backgrounds and environmental conditions.

Together, this study establishes a conceptual and computational framework for transforming regional haplotypes into practical breeding units. By enabling rational parental selection and precise pyramiding of favorable genetic combinations, HAPBDB provides a foundation for next-generation precision breeding strategies in crops.

## Materials and methods

### The construction of lettuce genomic panel

To construct a comprehensive and highly representative genomic resource for lettuce, we collected publicly available variant call format (VCF) datasets from 631 lettuce accessions, including RNA-seq-derived variants from 240 accessions of a natural population ^31^ and whole-genome resequencing (WGS)-derived variants from 396 accessions of a MAGIC population ^12^. All variants were called against the Lactuca sativa cv. Salinas reference genome (version 8). According to the original studies, RNA-seq reads were aligned using HISAT2, whereas WGS reads were aligned using BWA-MEM prior to variant calling. The downloaded variant datasets comprised 401,272 SNPs from the RNA-seq dataset and 46,217,729 SNPs from the MAGIC population WGS dataset. To generate a unified genotype matrix across different sequencing methodologies, we used VCFtools (version 0.1.16) to intersect and integrate the SNPs, resulting in a final dataset comprising 229,878 high-quality SNPs across 631 accessions, referred to as the LG breeding panel, which was used for subsequent haplotype analysis.

### Haplotype analysis based on HAPBDB

To delineate and visualize haplotype structures from population-scale genotype data, we developed a computational pipeline integrating both strict and hierarchical clustering strategies implemented in HAPBDB (**Fig. 6**). The workflow initiates with a VCF file containing genotypic information from multiple accessions. From these raw data, two genotype matrices are generated from these data: a binary-encoded matrix representing genotype states for clustering analyses and a nucleotide-level genotype matrix (A/T/C/G) for downstream haplotype visualization.

**Figure 6.**
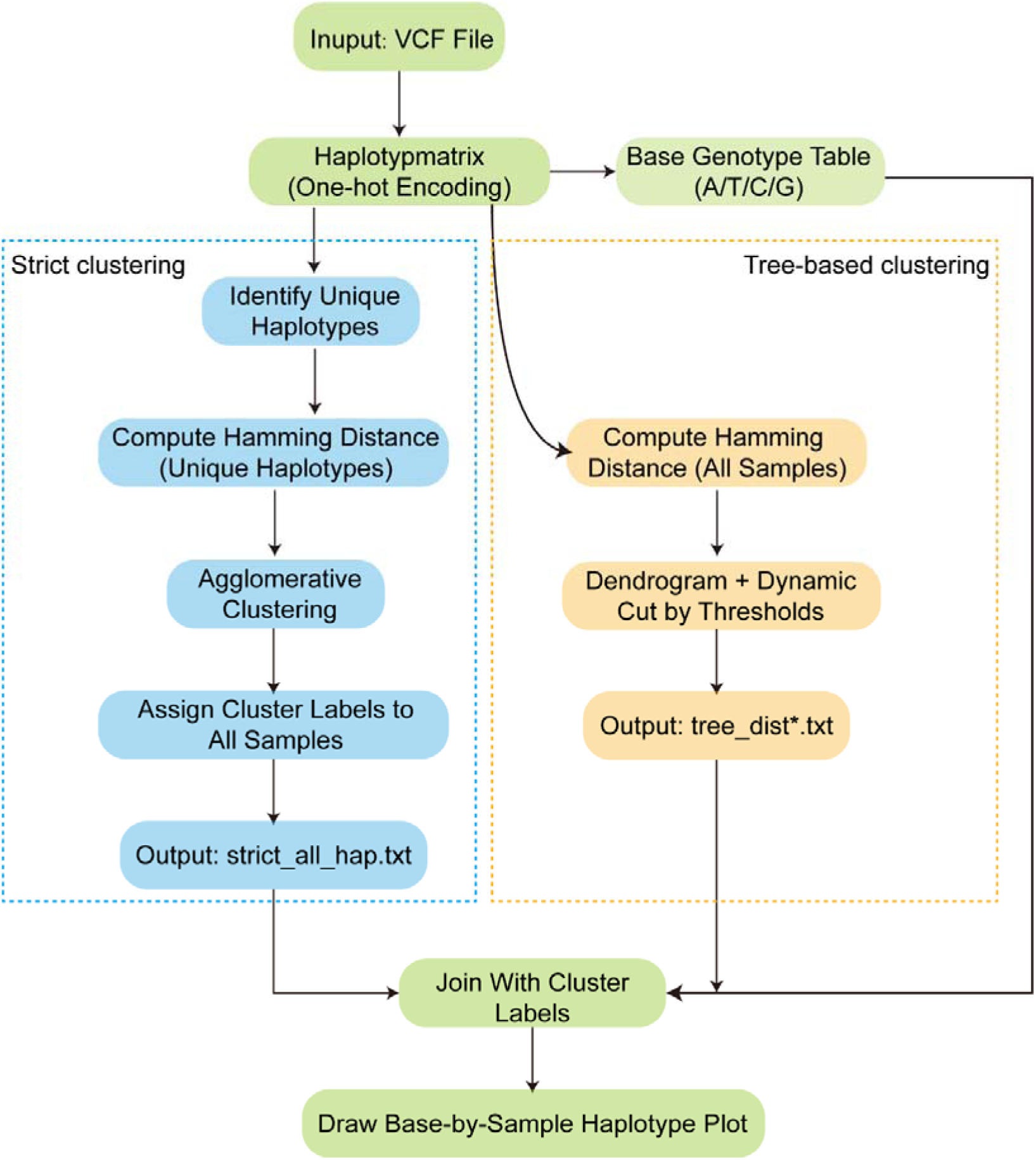
Computational pipelines implemented in HAPBDB.

For strict haplotype clustering module, unique genotype patterns were first identified from the binary-encoded genotype matrix and accessions sharing identical genotype configurations were assigned to a single consensus haplotype. Following this primary classification, pairwise Hamming distances among these representative haplotypes were subsequently calculated and subjected to visualize their genetic relationships by hierarchical clustering.

In parallel, HAPBDB performed a tree-guided clustering approach based on the pairwise hamming distances across all accessions using an average-linkage algorithm. To define biologically meaningful haplotypes, the dendrogram is iteratively partitioned using decreasing distance thresholds until two predefined reference accessions (typically representing favorable and unfavorable haplotypes) are assigned into distinct clusters. The resulting cluster assignments are subsequently used as the final haplotype classification for downstream genetic and breeding analyses.

For haplotype visualization, the nucleotide-level genotype matrix is integrated with haplotype cluster labels derived from either clustering strategy. For each haplotype, a representative accession is selected to display nucleotide variation across polymorphic sites.

To visualize the relationships among haplotypes, one representative accession was selected from each haplotype group identified by HAPBDB pipeline. The pairwise genetic distances among representative accessions were extracted from the complete distance matrix generated in the clustering phase. Hierarchical clustering was performed using the average-linkage method implemented in the SciPy Python package. The dendrogram was generated using the dendrogram function and visualized with Matplotlib in Python. To enhance the informative value of the plot, the total number of accessions within each haplotype was explicitly annotated adjacent to its corresponding haplotype label on the tree.

Ultimately, HAPBDB generates a series of graphical outputs, including haplotype dendrograms, Hamming distance heatmaps, and haplotype-by-site nucleotide matrices, providing intuitive visualization of haplotype diversity and genetic relationships within target genomic regions as well as a summary report.

### Multi-locus elite haplotype aggregation analysis

To identify accessions carrying complementary favorable haplotypes across multiple QTL loci, accessions harboring elite haplotypes at each specific target locus were extracted from the HAPBDB results and grouped into distinct accession sets. To map the co-occurrence of these elite loci, pairwise and higher-order intersections among these sets were calculated using custom Python scripts based on set operations. The distribution of shared and unique accessions among favorable haplotype groups were visualized using UpSet plots implemented in the Python packages, respectively. The resulting overlap patterns were subsequently used to prioritize parental accessions for favorable haplotype pyramiding.

### Plant materials and hybrid

All selected lettuce accessions were cultivated in a controlled greenhouse environment at Huazhong Agricultural University (Wuhan, China). Plants were grown under standard irrigation and fertilization conditions and underwent bolting and flowering within 3-4 months. Controlled crosses were conducted according to predefined hybrid combinations, with emasculation and pollination performed around sunrise. The hybrid seeds were harvested 10-15 days post-pollination when seeds reached maturity. The resulting F hybrids, along with parental lines, were subsequently transplanted and grown under the same cultivation conditions for further analysis.

### DNA extraction and molecular marker validation

Leaf tissues from all plant materials used in this study, including parental lines and F hybrids, were collected for genomic DNA extraction. DNA was isolated using the CTAB method as previously described ^32^. The InDel markers capable of distinguishing the original QTL mapping parental lines were identified using the PSVGT toolkit, and corresponding PCR primers were designed for subsequent genotyping. All primer sequences are listed in **Table S5**. PCR amplification was performed using the designed primers, and the products were separated by electrophoresis on a 1% agarose gel. Haplotypes corresponding to different QTLs were identified based on band size variations visualized on the gels.

## Code availability

All code is available at https://github.com/lgbTime/HAPBDB under the MIT License.

## Acknowledgements

We thank the high-performance computing platform at the National Key Laboratory of Crop Genetic Improvement at Huazhong Agricultural University. This research was supported by grants from the National Key Research and Development Program of China (2023YFF1000100), the scientific research start-up funding from Hubei Hongshan Laboratory (award No. 11020102), the Fundamental Research Funds for the Central Universities (Program No.2662024SZ005), the earmarked fund for CARS (CARS-21-A13) and the Young Scientist Fostering Funds for the National Key Laboratory for Germplasm Innovation & Utilization of Horticultural Crops.

## Author contributions

X.W. conceived and managed this work. G.L designed the framework. G.L. and Z.T. implemented the framework. Z.T., L.X. J.Z. and M.W. performed data analysis and experiments. G.L., Z.T., and X.W. wrote the manuscript.

