## Supplemental Fig for "A haplotype-based breeding framework for the precise pyramiding of elite QTL alleles: a lettuce case study"

### Supplementary information

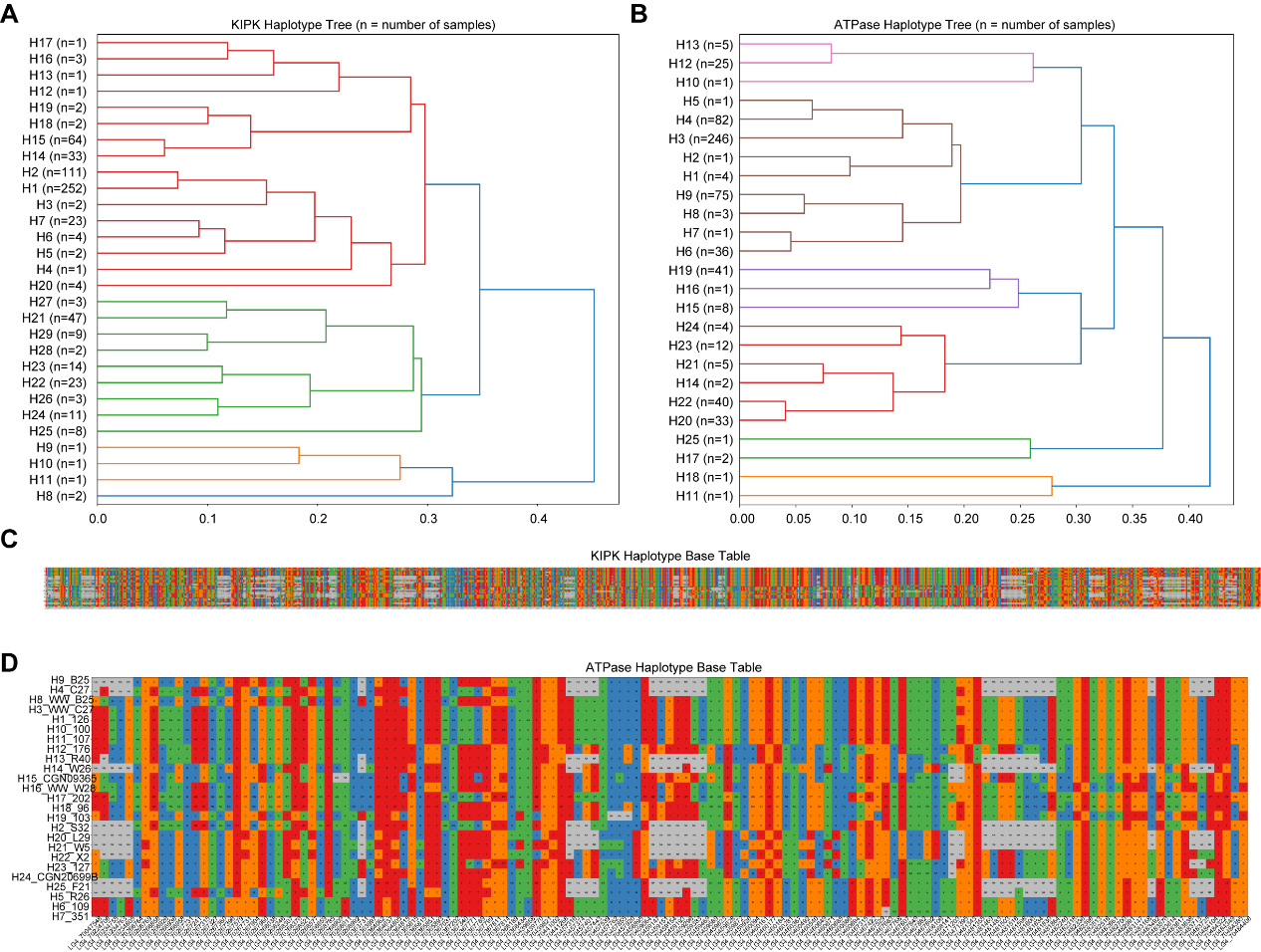

**Figure S1** **Regional haplotype dissection of the *LsKIPK* and *LsATPase* locus using HAPBDB.** (**A-B**) Dendrogram showing the phylogenetic relationships among *LsKIPK* and *LsATPase* regional haplotypes reconstructed by HAPBDB, with branch lengths proportional to pairwise Hamming distances. (**C-D**) Haplotype-genotype matrix of *LsKIPK* and *LsATPase*.

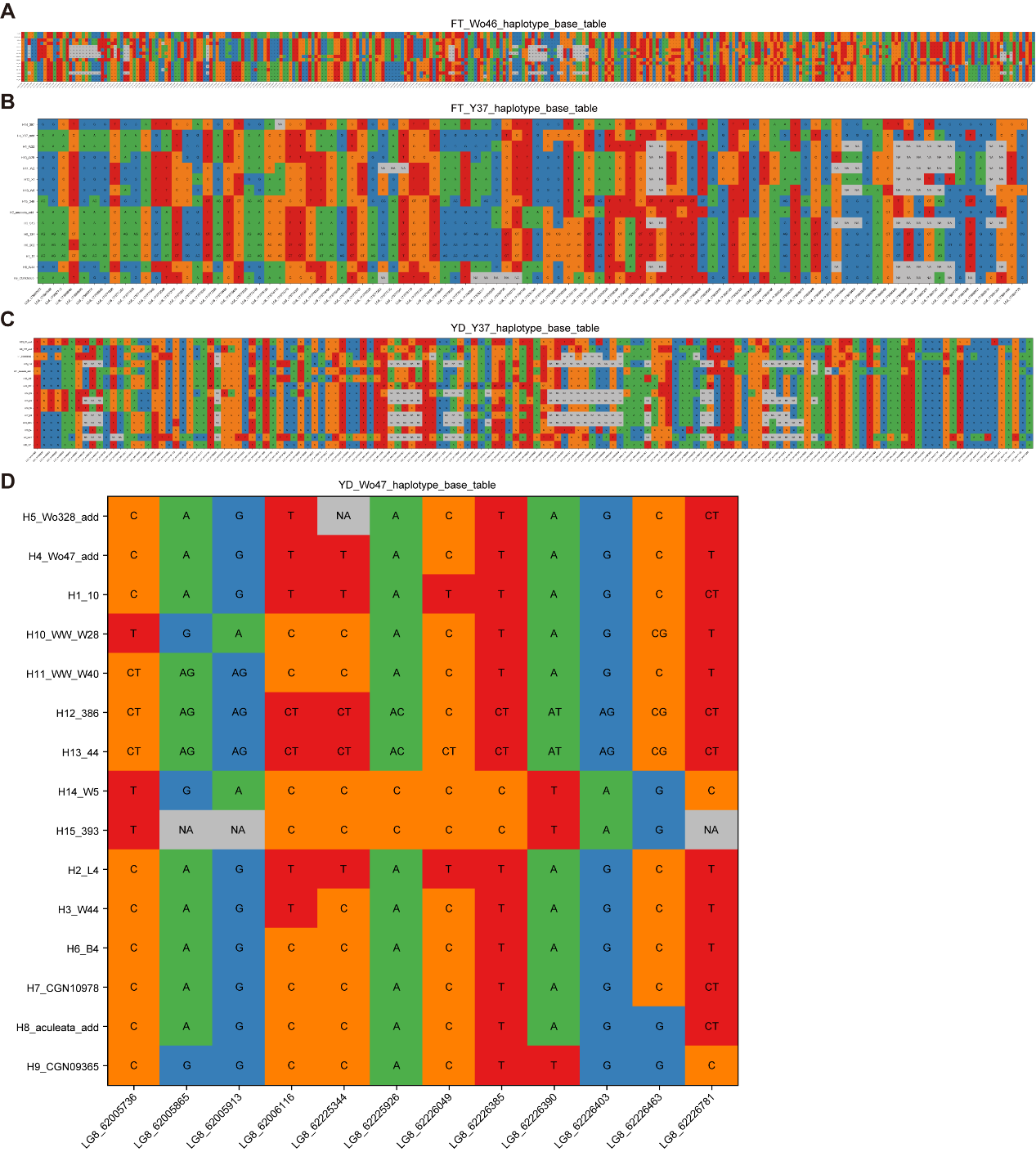

**Figure S2** **Genomic architecture and regional haplotype-genotype matrices.** (**A**) FT_Wo46. (**B**) FT_Y37. (**C**) YD_Y37. (**D**) YD_Wo47.

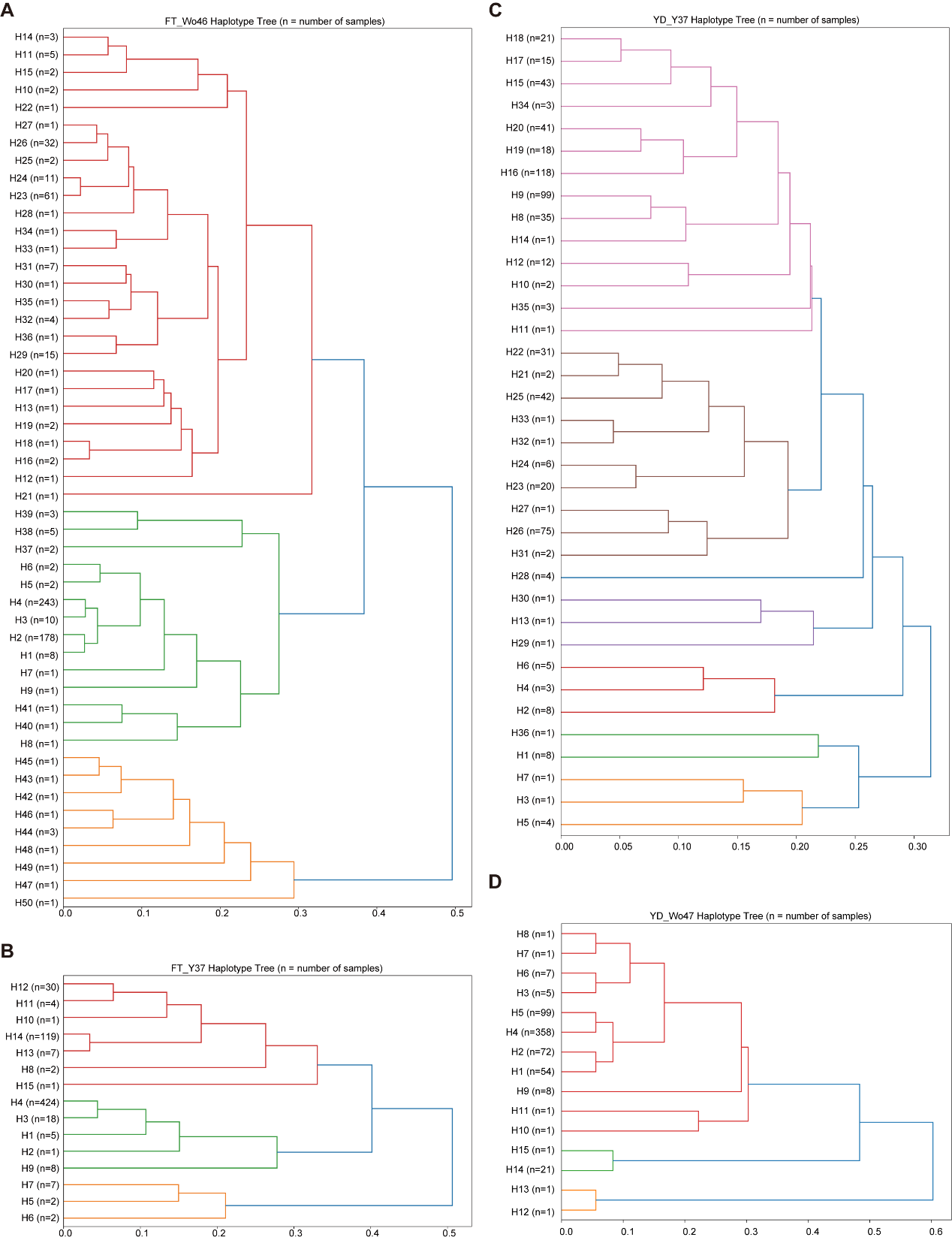

**Figure S3** **Phylogenetic dendrograms of regional haplotypes reconstructed by HAPBDB for four QTLs.** (**A**) FT_Wo46. (**B**) FT_Y37. (**C**) YD_Y37. (**D**) YD_Wo47. Branch lengths are proportional to pairwise Hamming distances among haplotypes. The vertical axis lists individual haplotypes, and the horizontal axis represents genetic distances. Closely clustered branches indicate higher sequence similarity, whereas longer branch lengths reflect greater genetic divergence.
